# Artery-on-chip demonstrates smooth muscle function comparable to both healthy and diseased living tissues

**DOI:** 10.1101/2025.03.06.641764

**Authors:** Danielle Yarbrough, Roy Chen, Jaron Shoemaker, Edwin Yu, Jennifer Arthur Ataam, Connor Amelung, Ioannis Karakikes, Sharon Gerecht

## Abstract

Arterial diseases affect the mechanical properties of blood vessels, which then alter their function via complex mechanisms. To develop and test effective treatments, microphysiological systems replicating the function and mechanics of a human artery are needed. Here, we establish an artery-on-chip (ARTOC) using vascular derivatives of human induced pluripotent stem cells (iPSCs) cultured with pulsatile flow on an electrospun fibrin hydrogel. ARTOCs have mature, laminated smooth muscle that expresses robust extracellular matrix and contractile proteins, contracts in response to intraluminal pressure and vasoagonists, and exhibits tissue mechanics comparable to those of human small-diameter arteries. Using real-time monitoring of radial distention and luminal pressure to inform computational fluid dynamics modeling, we show that we can effectively tune biomechanical cues using fibrin scaffold thickness and luminal flow rate. We successfully tune these cues to promote the survival and function of both endothelial and smooth muscle cells simultaneously in the ARTOC. To test the ARTOC as a disease modeling platform, we first use non-isogenic iPSC-derived smooth muscle cells from a polycythemia patient, and we find significantly altered cell phenotype and increased vessel wall stiffness compared to controls. We then test a novel isogenic disease model in ARTOCs from iPSCs CRISPR-edited with the LMNA Hutchinson-Guilford Progeria Syndrome (LMNA G608G; LMNA^HGPS^) mutation. LMNA^HGPS^ ARTOCs show extracellular matrix accumulation, medial layer loss, premature senescence, and loss of tissue elasticity and ductility. With this work, we establish the ARTOC as a platform for basic and translational studies of arterial diseases, bridging the current gap in linking protein expression and cell phenotype to tissue mechanics and function in small-diameter arteries.

## INTRODUCTION

Cardiovascular disease, and arterial disease in particular, is a leading cause of death globally.^1^ The root causes of life-threatening vascular disease range from environmental triggers to rare heritable genetic conditions, thus it is essential to identify the myriad molecular pathways that initiate vascular tissue degradation and dysfunction.^2^ While clinical studies are necessary for a better understanding of the pathology, they are often undertaken at a later stage of disease in which systemic dysfunction may be too far for certain treatments to halt or reverse disease progression.^3^

Microphsiological systems (MPS), miniaturized tissue models, can accelerate fundamental and translational studies to uncover key mechanisms of earlier vascular disease progression that cannot be observed in clinical settings.^4–8^ Toward the study of arterial disease, MPS should recapitulate a mature, multicellular, multilayered arterial tissue with smooth muscle that withstands and readily responds to physiologically relevant stimuli. Pioneering work in engineered arterial tissue focused on large-scale models for surgical applicability, and these systems were typically comprised of multiple vascular cell types cultured for extended periods of time, often integrated with strong synthetic scaffolds.^9–11^ Other approaches have since improved construct design and throughput by using flow-induced maturation, biomaterial innovations, and new cell seeding techniques such as bioprinting.^12–15^ Still, larger constructs can require anywhere from 1 to 3 months of culture to achieve mature, strong, and functional tissue.^16^

Advances in the usage of human iPSCs have provided a promising option for an accessible, renewable, and patient-specific cell source with applicability for disease modeling and precision medicine^17,18^. The proliferation capacity of iPSC-derived cells, their low batch-to-batch variability, and their genetic modifiability make them an excellent cell source to model blood vessels in MPS. Multicellular MPS seeded with iPSC-derived vascular cells have been generated successfully for both in vitro modeling and clinical applications, and many of them achieve similar anatomy to native vessels and demonstrate functional endothelium.^19–23^ Additionally, exposure of tissues to pulsatile luminal flow and strain has been shown to induce iPSC-derived smooth muscle cell (iSMC) maturation and mechanical strength prior to in vivo implantation.^24^

However, the main challenge in engineering arterial MPS in vitro has been in achieving a truly functional vascular smooth muscle tissue that responds readily to both pressure changes and vasoagonists, while enduring physiologically relevant pulsatile flow. Such functional vascular smooth muscle tissue could provide insights into disease progression and serve as a platform for testing therapeutics. The challenge in achieving this is partly because endothelial cells (ECs) remain the primary focus of most vascular MPS.^25,26^ Although ECs are essential for vascular signaling and responding to signals from the bloodstream, arterial tone is ultimately responsible for severe cardiovascular events or diseases.^27,28^ The arterial tone is dominated by the properties of both the SMCs and their endogenous extracellular matrix (ECM).^29–31^ The complex mechanisms that regulate arterial tone in vivo, and the difficulty in generating mature smooth muscle in vitro, have led to a comparatively limited understanding of the independent role SMCs play in arterial diseases, separate from endothelial signaling^32^. Additionally, the small artery vascular niche (less than 1 mm diameter) requires both high strength and elastic smooth muscle tissue in addition to a culture system operating on a microfluidic scale, and these constraints remain a challenge to achieve in vitro.^33,34^

Here, we develop an artery-on-chip (ARTOC) using iPSC-derived cells that recapitulate cellular organization and biomechanical properties of native human small arteries. We show distinctive, uniform, smooth muscle tissue with organized ECM that significantly improves ARTOC elasticity and drives pulsatile ARTOC contraction. Meanwhile, when ECs are incorporated into the ARTOC, they align in response to shear stress and contribute to barrier function. In a disease model from a polycythemia patient donor and an isogenic disease model for Hutchinson-Gilford Progeria Syndrome (HGPS), the ARTOC reproduces the clinically established tissue pathophysiology and abnormal SMC phenotypes. Thus, we show that a single ARTOC sample can connect clinically observed disease-specific arterial mechanics to discrete protein and transcript expression. With this important development for arterial disease modeling in vitro, we propose the ARTOC as a robust and versatile platform for studying early arterial disease progression and screening potential therapeutics for these diseases.

## RESULTS

### Smooth muscle cells successfully integrate with fibrin scaffold and modulate ARTOC tissue mechanics

To balance the need for mechanical strength and the goal of developing an anatomically accurate ARTOC, we used scaffolds composed of an electrospun fibrin hydrogel assembled into a tubular shape to generate aligned nanofibers that facilitate cell adherence and alignment ^35^ **(Fig. S1A)**. We generated iSMC-seeded ARTOCs to isolate the changes in bulk properties as a direct result of the iSMCs experiencing dynamic culture conditions. To assemble the ARTOC, iSMCs were cultured in a monolayer and physically rolled as a single cell sheet around the fibrin tubular construct (approximately 0.6 mm inner diameter) that was mounted on hollow cannulas in custom bioreactor chambers (**Fig. 1A**). Pulsatile flow via a peristaltic pump allowed for gas exchange through luminal media and induced circumferential strain to stimulate tissue maturation and biomechanical response (**Fig. 1B, Fig. S1E-F)**. Within 1 week of culture, live imaging of tissue cross sections showed that the addition of iSMCs significantly increased the total wall thickness in ARTOC by 2-fold (**Fig. 1C-D**) and significantly decreased the inner diameter through cell contraction around the fibrin scaffold (**Fig. 1E**), demonstrating the presence of an intact layer of contractile tissue functioning as a single unit. The resultant iSMC-populated ARTOC exhibited significantly increased burst pressure compared to fibrin-only constructs (**Fig. 1F**), withstanding a luminal pressure almost 3-fold higher than healthy physiological pressure.

**Figure 1.**
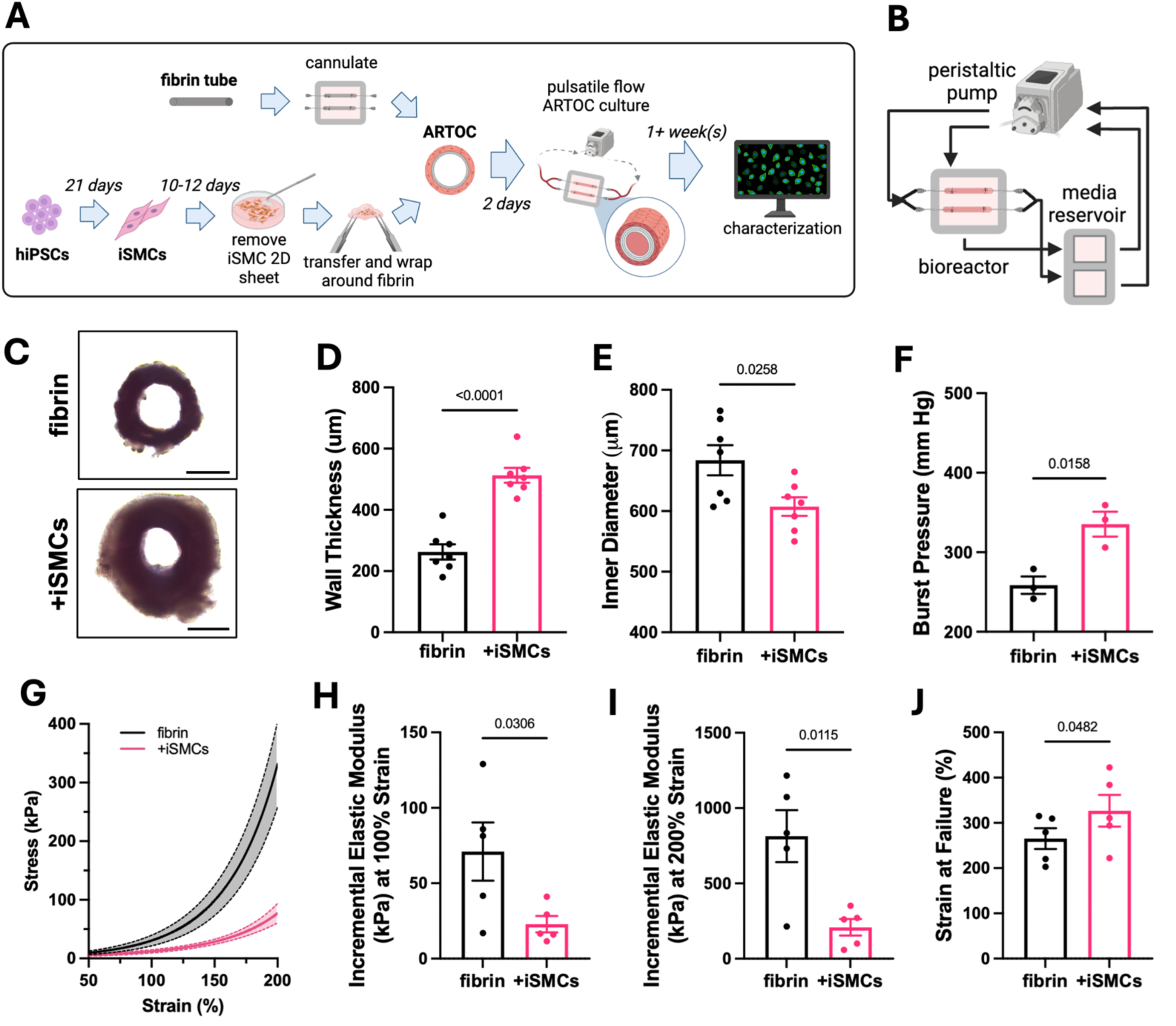
Artery-on-chip integration. **(A)** schematic of step-wise approach for iSMC derivation and ARTOC construction. **(B)** schematic of the setup of the pulsatile flow and bioreactor culture system. **(C)** widefield cross-sectional images of fibrin only or live iSMC-seeded ARTOC, scale=500um. **(D)** Total wall thickness, N=7, (n=2 each N) **(E)** inner diameter, N=7, (n=2 each N) and (**F)** Burst pressure for fibrin only (black) vs. iSMC-seeded (pink) ARTOC, N=3, (n=1-2 each N). **(G)** Stress v. strain curves for radial tensile testing of fibrin vs. iSMC-seeded ARTOC. Incremental elastic modulus at **(H)** low strain and at **(I)** high strain, and **(J)** strain at ultimate failure for fibrin-only v. iSMC-seeded ARTOC, N=5, (n=2-3 each N)

After 1 week of culture under pulsatile flow, constructs were subjected to radial tensile pulling until complete failure to generate stress-strain curves (**Fig. 1G**). Cellularized ARTOCs withstood higher strains (**Fig. 1H**) and exhibited decreased elastic moduli in both the low-strain (**Fig. 1I**) and high-strain (**Fig. 1J**) regions compared to fibrin-only constructs, indicating significantly increased elasticity due to the integrated iSMCs. While strain at failure was increased in cellularized ARTOC, ultimate tensile stress was significantly decreased compared to fibrin-only controls **(Fig S1B-D).** This may be due to the onset of fibrin degradation by adhered iSMCs, thus temporarily compromising ultimate ARTOC circumferential tensile strength but improving ARTOC elasticity.

These results show that the iSMC seeding method and ARTOC dynamic culture system successfully integrate living iSMC tissue with our tubular fibrin hydrogels. This integrated tissue-material construct significantly improves elasticity, ductility, and burst pressure compared to the material alone, achieving elastic moduli comparable to ex vivo porcine pulmonary artery sections of similar inner diameter (0.599 mm wall thickness, 50 kPa at low strain and 93 kPa high strain^36^) and slightly lower than human carotid arteries of larger inner diameter (7.7 mm ID, 0.61 mm wall thickness, 150 kPa at low strain and 750 kPa at high strain^37^). Taking into account differences in such in vivo testing methods, we can conclude that the ARTOC sufficiently mimics bulk mechanical properties of native small-diameter arteries. To our knowledge, this is a novel achievement for in vitro systems modeling this scale in the vascular niche (less than 1 mm diameter), and it demonstrates the robust nature of our ARTOC fabrication method.

### iSMC tissue reponse is promoted by tuning the pulsatile flow regime and fibrin construct fabrication

In the body, arteries readily respond to biomechanical cues as a necessity, but these cues are often not applied to cells in vitro, resulting in differences in cell phenotype and function compared to native cells. We subjected the ARTOC to pulsatile flow to apply biomechanical cues that induce cellular maturation and contractile response in our iSMCs. Through this, we aimed to achieve physiologically relevant levels of both radial distention and wall shear stress (WSS), which directly impact both the iSMCs and iECs present in the ARTOC.

Fibrin constructs were analyzed in situ for pulsatile distention and luminal pressure profile using a combination of optical micrometry and in-line pressure sensors to determine a range of physiologically relevant flow rates for the iSMCs (Fig. 2A). We fabricated thin and thick fibrin tubular constructs and found that systolic and diastolic internal pressure increased linearly with luminal flow rate, and physiologically relevant pressures of approximately 120/65 mm Hg at a flow rate of 0.8 mL/min corresponded to 1.2% or 0.7% distention in thin or thick fibrin constructs, respectively (Fig. 2B). The circumferential distention experienced by iSMCs was further tunable by changing the fibrin scaffold thickness and luminal fluid flow rate (**Fig. 2C-D**). Importantly, iSMCs appeared to contribute to the distention under flow, increasing from approximately 1% in fibrin constructs up to 5% in cellularized ARTOCs at high flow rates (**Fig. 2E**).

**Figure 2.**
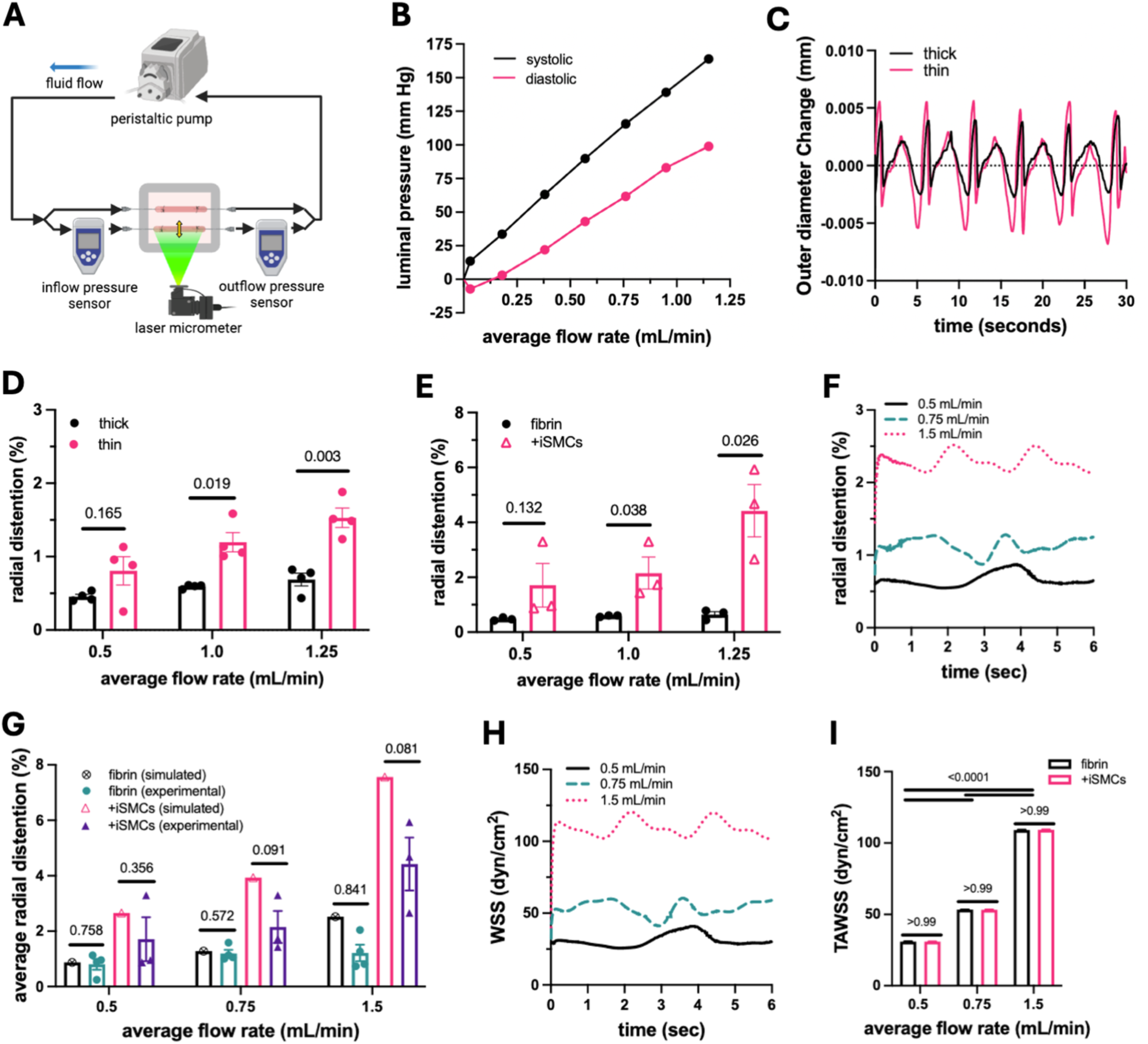
Pulsatile flow regime characterization. **(A)** Schematic for in situ measurement of internal pressure (inflow and outflow pressure sensors) and outer diameter distention (laser micrometer) profiles with time. **(B)** Internal systolic and diastolic pressure in fibrin only scaffolds in bioreactors measured at varying flow rates. **(C)** Outer diameter readout over 30 seconds in thick or thin fibrin-only scaffolds. **(D)** In situ percent radial distention for thin or thick fibrin scaffolds at varying flow rates, N=4 (n=2 each N). **(E)** Percent radial distention in fibrin vs. ARTOC at varying flow rates, N=3 (n=2 each N). **(F)** CFD-modeled circumferential distention in fibrin scaffolds at varying flow rates over 6 seconds. **(G)** simulated (open symbols) vs. experimental (solid symbols) circumferential distention for fibrin scaffolds and ARTOC at varying flow rates, N=3 (n=2 each N). **(H)** CFD-modeled WSS over 6 seconds for fibrin constructs at varying flow rates. **(I)** Simulated TAWSS over 1 minute for fibrin scaffolds and ARTOC at varying flow rates, N=3, (n=2 each N).

To determine a physiologically relevant flow rate that would be compatible with the luminal endothelization in addition to the externally seeded iSMCs, we initially estimated the average wall shear stress experienced in fibrin constructs corresponding to our experimental mean flow rate of 0.5 mL/min to be 4 dyn/cm^2^ **(Fig. S2A)** using a simplified Hagen-Pouseille flow assumption.^38^ Because we apply pulsatile flow to an elastic material in our system, we hypothesized that Pouseille flow estimation would not be sufficient to estimate the wall shear stress applied to endothelial cells, and we sought to model it more accurately in real-time. To accomplish this, we used in-line pressure measurements to fit an oscillatory flow model for an elastic hollow cylinder with open-source SimVascular computational fluid dynamics (CFD) software^39^ **(Fig. S2B-C; Video S1-2)**. ARTOC distention was modeled first to confirm that the CFD simulation sufficiently matched our experimentally measured circumferential distention (**Fig. 2F**). Indeed, our experimental distention values were not significantly different from the simulation’s expected distention values for any modeled flow rates in fibrin constructs or cellularized ARTOCs (**Fig. 2G**). These findings show that the CFD simulation is capable of accurately predicting other flow parameters for our system, such as WSS. At a flow rate of 0.5 mL/min, the time-averaged wall shear stress (TAWSS) approximated 30 dyn/cm^2^ with a maximum wall shear stress of 41 dyn/cm^2^ at peak systolic velocity (**Fig. 2H**), which was 10-fold higher than the initial Hagen-Pouseille estimation. Further, the TAWSS significantly increased with increasing flow rates but did not vary significantly between fibrin and ARTOC simulations (**Fig. 2I**).

From these results, we conclude that the easily tunable wall thickness of the fibrin constructs induced a desired circumferential strain at varying luminal flow rates, directly acting as a biomechanical cue to the iSMCs seeded on the fibrin construct. The iSMCs also contributed to additional circumferential distention and contraction under pulsatile flow, showing that ARTOCs are functionally mature and exhibit whole-tissue contractility in response to a biomechanical stimulus. Lastly, we used in situ luminal pressure and distention measurements to optimize an oscillatory flow CFD model and accurately predict instantaneous WSS at varying flow rates in ARTOC. The model parameters were successfully optimized for both fibrin and ARTOCs to choose an appropriate flow rate for applying physiologically relevant WSS for the iECs and circumferential distention for the iSMCs. These results underscore the importance of using thorough CFD to accurately estimate the mechanical forces being applied to cells in vascular MPS that incorporate flow. Concurrently, the dual in situ measurement of diameter and pressure under pulsatile flow highlights the dynamic nature of the ARTOCs and their capability to readily respond to mechanical stimuli as arteries do in the human body.

### Co-cultured ARTOCs recapitulate healthy mature and functional arterial tissue

While SMCs exhibit the contractile function of arteries, the ECs lining the lumen of blood vessels regulate response to signals from the bloodstream. Thus, a fully functional ARTOC must be capable of including ECs in co-culture with SMCs to examine vessel wall permeability and EC-mediated response to stimuli. To this end, fibrin tubes were seeded luminally with a high density of isogenic iECs **(Fig. S3A**). Within 3 days, iECs formed a confluent monolayer with vascular endothelial cadherin (VE-cad) expression localized at the cell-cell junctions, demonstrating an intact endothelial barrier (**Fig. 3A**). After iSMCs were added to the external surface of the fibrin constructs and the co-culture ARTOC were subjected to luminal flow for at least 1 day, both cell types retained their phenotype and continued to express cell-specific markers: VE-cad for iECs and smooth muscle myosin heavy chain (SM-MHC) for iSMCs, with iECs aligning parallel to flow (**Fig. 3B, Fig. S3B)**. Endothelial barrier integrity was also validated by measuring the extravasation of FITC-Dextran (70kDa) from the luminal media into the external media chamber while under flow for 1 day. ARTOCs with iECs showed decreased permeability compared to fibrin alone and at comparable levels to published vessel wall permeability coefficients,^40^ indicating a functional endothelial barrier (**Fig. 3C**).

**Figure 3.**
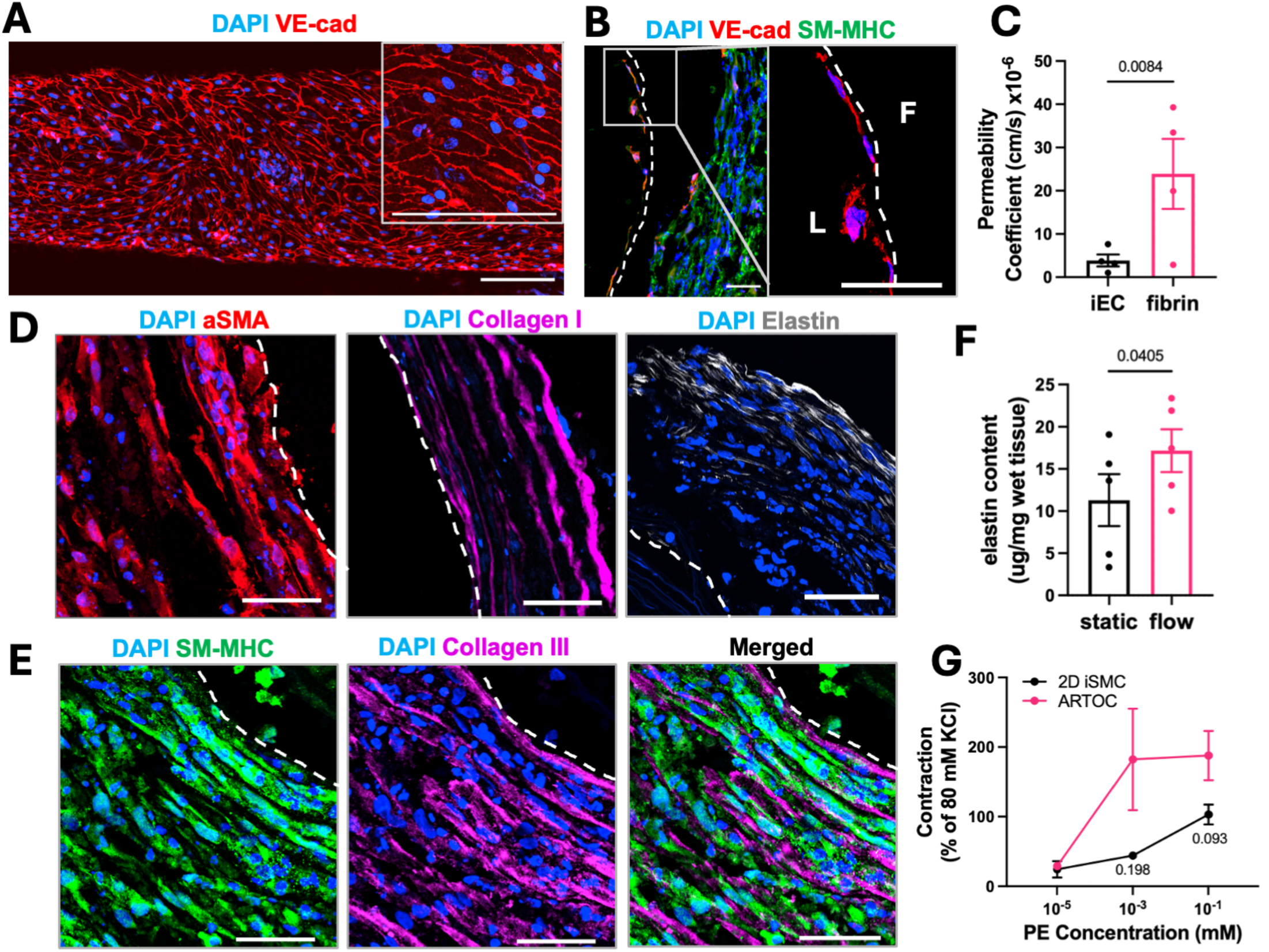
Co-culture ARTOC phenotypic maturity and functionality. **(A)** En face staining of iEC-seeded ARTOC lumen for VE-cad (red) after 3 days of static culture in bioreactors, scale=150um **(B)** Cross-section stain showing iECs and iSMCs forming a co-culture ARTOC expressing VE-cad (red) and SM-MHC (green), insert showing iECs in lumen, L=lumen, F=fibrin, scale=20 um. **(C)** Calculated permeability coefficient for iEC-seeded or fibrin scaffolds after perfusion of 70 kDa FITC-Dextran, N=5. **(D)** Cross-section stains for SMC markers alpha-SMA (red), collagen I (magenta), and elastin (white), and **(E)** co-staining for SM-MHC (green) and collagen III (magenta), dotted line=lumen, scale=50 um. **(F)** Fastin Elastin Assay quantification of mature elastin content in ARTOC cultured for 1 week in static or flow conditions, N=5 (n=2 each N). **(G)** Percent contraction in ARTOC cross-sectional area vs. 2D iSMC area in response to increasing PE doses as a fraction of maximum KCl-induced contraction, N=3 (n=2 each).

After at least 1 week of dynamic culture in the bioreactors, iSMCs formed a dense, multilayered muscle tissue that expressed contractile markers α-smooth muscle actin (aSMA) and SM-MHC, laminated collagen I and collagen III, and mature elastin fibers (**Fig. 3D-E**). Biochemical quantification indicated that elastin content was significantly increased by 1 week of culture with pulsatile flow, and approximated half of that in native ex vivo veins and arteries from murine and porcine models, respectively^41,42^(**Fig. 3F**). Additionally, iSMC contractile function was validated by measuring area contraction after sequential treatments with vasoconstrictors potassium chloride (KCl) at 80 mM and phenylephrine (PE) at 10^-5^-10^-1^ mM. We found that iSMC-only ARTOC cross-sectional area contracted in a dose-responsive manner and to a slightly greater extent than monolayer iSMCs after PE treatment (as a fraction of KCl-induced contraction) and in a similar manner to ex vivo arteries from human patients^43^ and animal models^44,45^ (**Fig. 3G**). These combined results demonstrate that the ARTOC is a phenotypically mature and functional in vitro arterial model that responds readily to vasoagonists and maintains barrier function under flow when iECs are incorporated. Further, the functional capabilities of both iECs and iSMCs in the ARTOC indicate a robust pharmacological response that could be exploited for pre-clinical screening of therapeutics.

### HIF2A mutated ARTOCs reflect cellular phenotype and polycythemia clinical indication

The ARTOC culture system is particularly useful as a platform to model vascular diseases that disproportionately affect human smooth muscle tissue. We thus sought to validate that the ARTOC can capture both abnormal cell phehotype and clinically observed tissue dysfunction in smooth muscle-related arterial disease. Patients with primary polycythemia often suffer from a variety of cardiovascular complications including hypertension and thromboses leading to strokes, and current standard treatment is focused on directly removing red blood cells through repeated venesection.^46–48^ This form of polycythemia can be induced by a heritable genetic mutation, one of which is a gain-of-function mutation in the hypoxia-inducible factor 2α (HIF2A) gene.^49^ In previous studies, we found phenotypic alterations in HIF2A-SMCs compared to wild-type (WT) SMCs derived from iPSCs, and HIF2A-GOF mice exhibited hypertension and increased arterial stiffness.^50^ Here, we sought to examine if these changes persist in the ARTOC.

We found that the HIF2A ARTOCs had increased protein expression of aSMA, acetylated α tubulin (ac-a-tubulin), and F-actin compared to WT ARTOCs. In contrast, collagen I and SM-MHC protein expression increased only slightly in an inconsistent manner (**Fig. 4A-B**). We next aimed to quantify viscoelasticity in the ARTOCs to determine if HIF2A ARTOCs recapitulated the arterial wall stiffness observed previously.^51^ While traditional viscoelastic measurements, such as rheology, adequately capture macro-scale viscoelastic elements, we aimed to better quantify the heterogeneity present at the cell scale using atomic force microscopy (AFM) with an oscillatory microrheology module. As opposed to traditional AFM that captures only the elastic Young’s Modulus (E), microrheology enables understanding of the viscoelastic components, such as the storage modulus (E’) and loss modulus (E’’) by oscillating at a known frequency.^52^ When microrheology was measured, HIF2A ARTOCs had both increased storage and loss modulus compared to WT at a 1 Hz indentation frequency (**Fig. 4C-D**), which indicates both increased tissue stiffness and viscosity, respectively, in the HIF2A ARTOCs. This difference in storage modulus was maintained across a full frequency sweep from 1 to 100 Hertz, while loss modulus difference began to narrow at higher frequencies (**Fig. 4E-F**). Meanwhile, the loss factor between storage and loss modulus remained similar between HIF2A and WT ARTOCs across all frequencies **(Fig. S4A)**. These results indicate that while the elastic stresses (E’) dominate the overall increase in HIF2A ARTOC stiffness regardless of deformation rate, the highest frequencies reveal more contribution by the viscous stresses (E”) in WT ARTOCs. This could be due to the observed actin overexpression in HIF2A iSMCs, keeping the cytoskeleton more intact and preventing the diseased ARTOCs from taking on a more viscous behavior at the high frequencies.^53^ With these results, we conclude that ARTOCs can not only capture the disease phenotype previously established in HIF2A-iSMCs, but they can also reproduce overall pathological tissue stiffness associated with polycythemia in the clinic.

**Figure 4.**
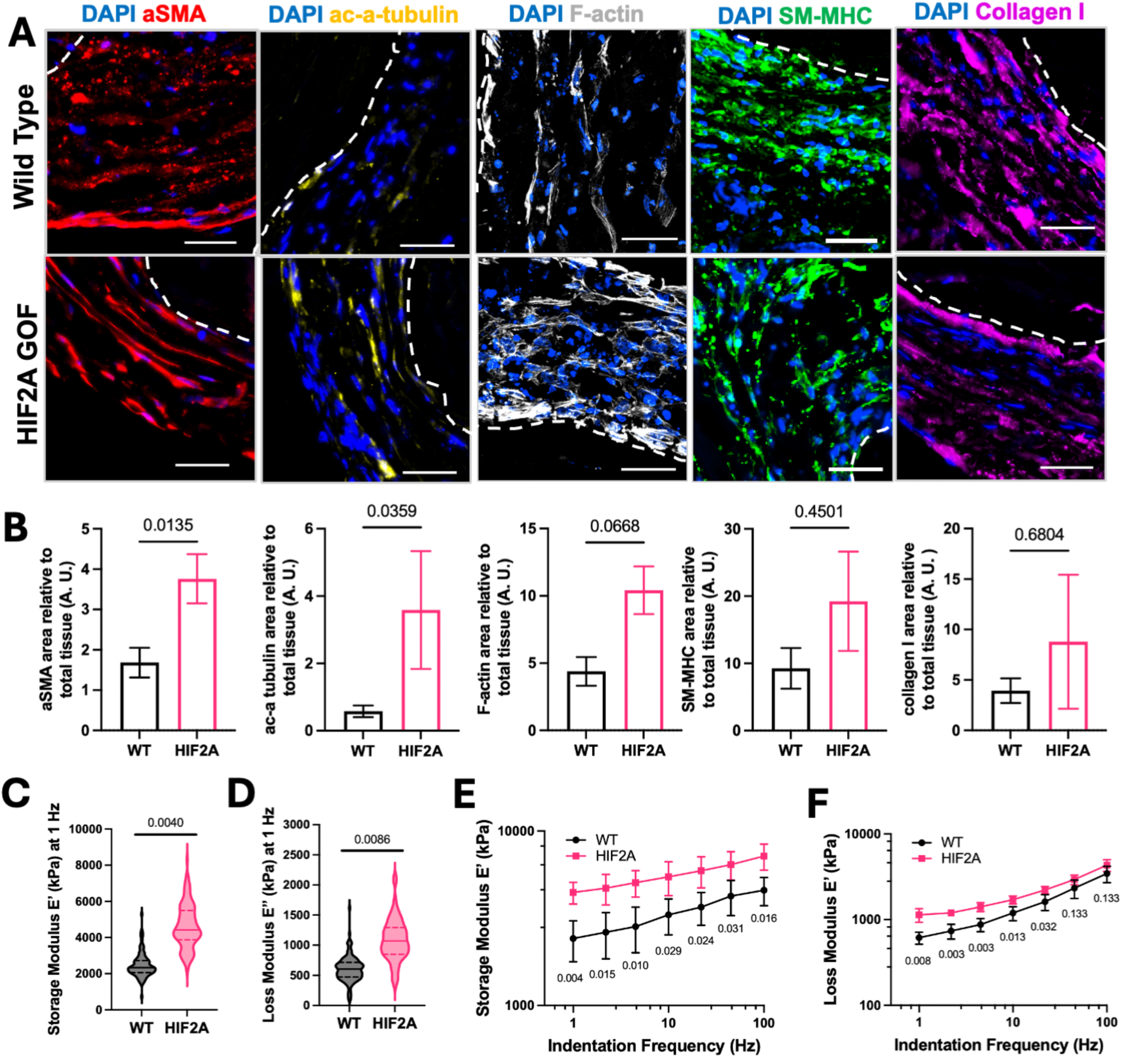
Cellular characteristics of HIF2A-GOF iSMCs are retained in ARTOCs. **(A)** Cross sections of wild type (top) and HIF2A (bottom) ARTOC stained for aSMA (red), ac-a-tubulin (yellow), F-actin (white), SM-MHC (green), and collagen I (magenta), dotted line=lumen, scale=50 um. **(B)** corresponding quantification of expression area normalized to total tissue for the same proteins, N=3 (n=5 images each N) **(C)** Storage modulus and **(D)** loss modulus at 1Hz in HIF2A or WT smooth muscle tissue as measured via AFM-microrheology, N=4 (n=65-130 each N). **(E)** Storage and **(F)** loss modulus in HIF2A or WT ARTOC measured over an entire 1 to 100 Hz frequency sweep, N=4 (n=65-130 each N).

### ARTOC recapitulates key aspects of HGPS smooth muscle tissue dysfunction

Human iPSC-based models enable personalized medicine applications and are particularly useful in studying rare genetic diseases. One such disease, Hutchinson Guilford Progeria Syndrome (HGPS) is often caused by a *de novo* mutation in the lamin A (LMNA) gene. This mutation leads leads to progerin accumulation which alters nuclear structure and function, causing premature senescence.^54^ The vascular cells are particularly affected by this pathology, so cardiovascular events such as stroke or severe atherosclerosis are the primary cause of premature death in patients.^55–57^ Because SMCs play a pivotal role in cardiovascular complications in HGPS patients, we hypothesize that the ARTOC can sufficiently model HGPS in vitro.^58^

We introduced LMNA HGPS mutation (c. 1824 C>T; p.Gly608Gly) at exon 11 in LMNA^WT^ iPSC line using cytosine base editing approach (CBE)^59^ (**Fig. 5A**). LMNA^HGPS^ iSMCs exhibited hallmarks of HGPS, including reduced proliferation rate (**Fig. 5B**), protein expression of progerin and decreased expression of lamin A (**Fig. 5C**), and typical nuclear envelope blebbing (**Fig.5D**). The LMNA^HGPS^ iSMCs were integrated into ARTOC, and after 1 week, they exhibited increased expression of collagen I, fibronectin, and senescence marker p21 when compared to LMNA^WT^ ARTOCs (**Fig. 5E-F**), alongside nuclear progerin expression and significant SMC loss **(Fig. S5A-B)** as previously seen in other vitro MPS for HGPS.^60^ Radial tensile testing showed that after an extended culture time of 3 weeks under flow, LMNA^HGPS^ ARTOCs had an increased elastic modulus at low and high strain as well as a slightly decreased strain at failure, indicating that they are less elastic and more brittle than LMNA^WT^ ARTOC (**Fig. 5G-J, Fig. S5C)**. These findings are consistent with clinical indications in which HGPS patient arteries exhibit excess ECM deposition and fibrosis, leading to arterial stiffness and the loss of vascular tone and the medial layer of SMCs.^61^ Altogether, these results show that our ARTOC can reproduce the HGPS phenotype and arterial pathology indicators, making them a viable model for studying HGPS-induced arterial dysfunction in vitro. Moreover, our use of iPSC-derived isogenic cells enables us to confirm the phenotype and bulk tissue pathology in LMNA^HGPS^ ARTOCs as a direct result of the induced lamin A mutation, independent of biological variability, thereby providing a test case for the ARTOC in precision medicine applications.

**Figure 5.**
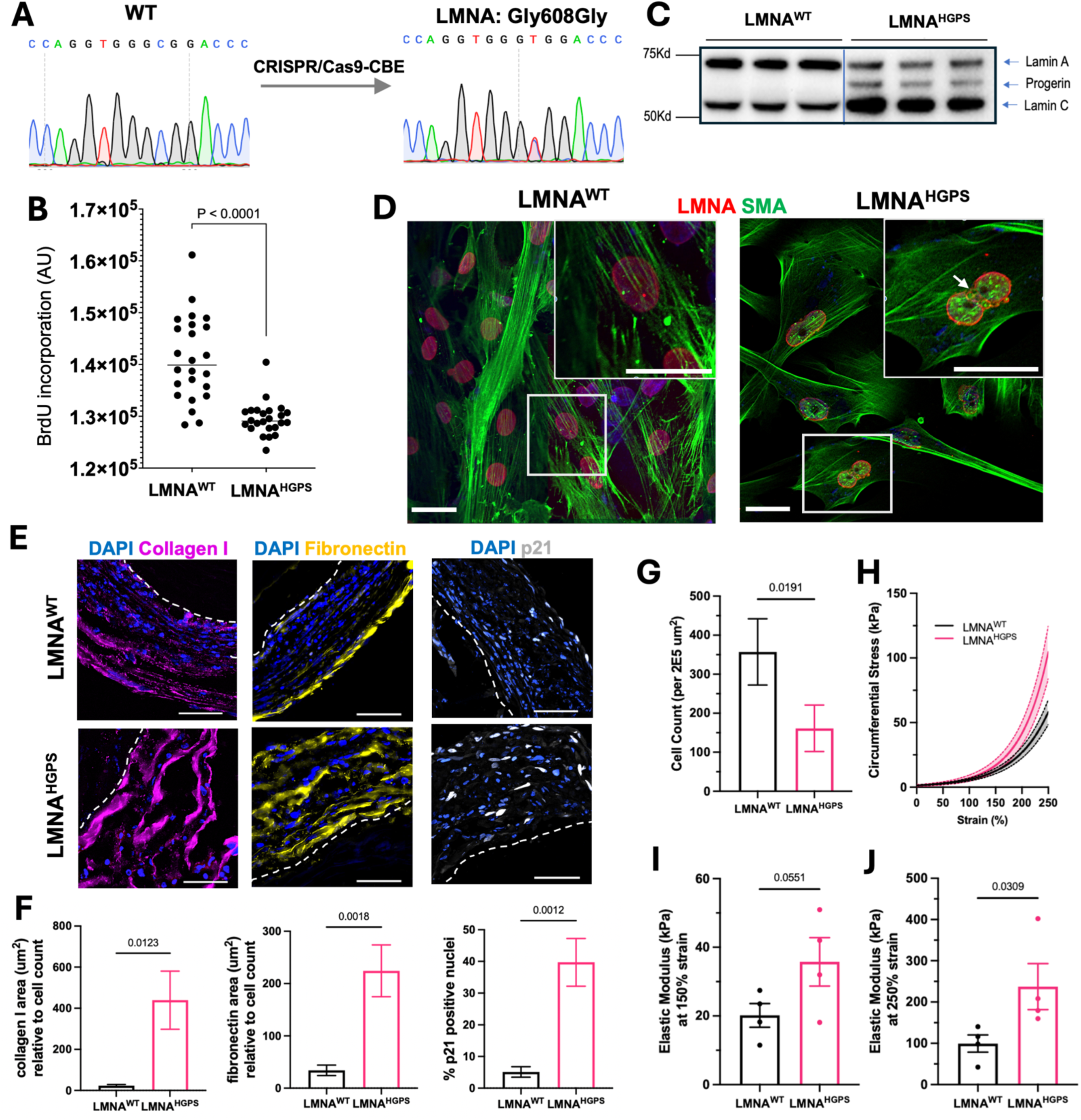
LMNA^HGPS^ ARTOC reproduces disease phenotype and in vivo tissue dysfunction. **(A)** iPSC and gene correction chromatogram**. (B)** Proliferation and **(C)** WB of Progerin, Lamina A and C of iSMCs derived from LMNA^WT^ and LMNA^HGPS^. **(D)** Immunostaining of isogenic iSMC carrying the *LMNA HGPS* mutation showing LMNA (red), SMA (green), and white arrow indicates blebbing nuclei, scale=50 um **(E)** Cross-sections of LMNA^WT^ (top) and LMNA^HGPS^(bottom) ARTOCs stained for collagen I (magenta), fibronectin (yellow), and p21 (white), dotted line=lumen, scale=50 um. **(F)** quantification of expression area normalized to cell number for the same three proteins, N=3 (n=5 images each N). **(G)** cell number in LMNA^WT^and LMNA^HGPS^, N=3 (n=5 images each N). Radial tensile testing of LMNA^WT^and LMNA^HGPS^ ARTOC cultured under flow for 3 weeks: **(H)** fitted stress v. strain curves, **(I)** incremental elastic modulus at 150% strain and **(J)** at 250% strain, N=4.

## DISCUSSION

Although the function of the smooth muscle is impaired in arterial diseases, their individual mechanistic role remains uncertain in many of these diseases due to the difficulty of in vitro modeling of human smooth muscle tissue.^62^ Despite significant advances, current in vitro arterial models lack a focus on fully characterizing a vasoactive smooth muscle layer that accurately mimics mature tissue structure, strength, and elasticity.^63^ In this work, we developed and characterized an ARTOC incorporating iSMCs, instructive hydrogel scaffold, and mechanical cues, resulting in rapidly matured and functional arterial tissue that effectively models multiple arterial diseases.

Advances in both smooth muscle seeding techniques, e.g. bioprinting, gel casting, and cell sheet wrapping^22^, and the increased use of iPSCs as a cell source, have yielded promising new in vitro arterial models, but rapid maturation (in less than 4 weeks) and in situ contractile function in these models remains elusive.^24,64,65^ We adapted a cell sheet wrapping approach and showed that within 9 days of total culture time on the construct, the multi-layered iSMCs integrated with and actively contracted around the fibrin scaffolds, significantly improving the elasticity, ductility, and burst pressure of the ARTOC, reaching comparable elastic moduli to large animal and human arteries (distal pulmonary and carotid, respectively).^36,37^ The iSMCs in the ARTOCs express mature contractile markers SM-MHC and aSMA, as well as the laminated collagen I and III fibers typical for arterioles and small arteries. Elastin appeared in a mature fibrillar structure, and elastin production increased under flow to approximately half that of large animal arteries in a relatively short time period.^42,66,67^ Finally, the smooth muscle tissue exhibited active contractility in response to circumferential strain and pharmacological agents, thus achieving a major functional benchmark.^44,68^ These results establish the ARTOC as a mature and functional model of human small arteries. The ARTOC not only matures comparatively much more quickly than the majority of in vitro artery models, but it is preeminent in achieving physiologically relevant arterial tissue mechanics at such a small scale (less than 1 mm diameter).

While this work focuses on recapitulating the arterial smooth muscle, iECs were also incorporated to demonstrate the feasibility of co-culture studies in the ARTOC. Using in situ pressure and distention measurements, we accurately simulated real-time WSS at various flow rates, confirming that it was largely unaffected by changes in overall ARTOC material properties. We could thus tune the flow rate to simultaneously achieve a desirable WSS for the iECs and appropriate radial distention for the iSMCs. Incorporated iECs formed a confluent luminal monolayer, displaying barrier function and alignment under flow and expressing VE-cad at cell junctions. Thus, a co-culture ARTOC as shown here, can be utilized to investigate EC-driven disease mechanisms and cell-cell communication across diverse contexts.

The ARTOC is particularly equipped to model diseases heavily affecting SMCs, and we were able to recapitulate two distinct diseases in both cellular phenotype and bulk tissue dysfunction similar to what is seen in patients. First, we modeled primary polycythemia induced by a HIF2A GOF mutation, using iPSCs from polycythemia patients or healthy donors. Previously, we have shown that this mutation alone resulted in aberrant expression of contractile and ECM proteins in iSMCs, and a HIF2A GOF mutation in a rodent model caused increased arterial stiffness and hypertension^50^. Our HIF2A ARTOCs also showed this aberrant phenotype, exhibiting tissue-wide increased stiffness and viscosity in addition to increased contractile protein expression. Next, we used an isogenic iPSC line from a healthy donor and induced a LMNA mutation to promote onset of the HGPS phenotype, characterized by progerin accumulation, cellular senescence, and nuclear blebbing. The LMNA^HGPS^ ARTOCs also exhibited senescence onset, massive loss of nuclei in the smooth muscle tissue, and excessive ECM accumulation, resulting in increased stiffness and slight loss of ductility.^69,70^ In both disease models, we linked protein expression to increased smooth muscle tissue dysfunction consistent with other in vitro and in vivo models,^60^ showing that the ARTOC is a versatile and robust in vitro platform capable of linking SMC mechanistic dysregulation to tissue-scale clinical indications in arterial diseases driven by SMCs.

In summary, we established a novel ARTOC that facilitates multiplexed analysis of both tissue-level arterial dysfunction and protein and transcript quantification for mechanistic studies. The ARTOC provides an important translational bridge between basic science studies conducted in vitro and complex pre-clinical animal models predicting human arterial responses. The ARTOC also fills a current gap in the ability to study human smooth muscle function in the small-diameter artery niche. In this work, we focus solely on the direct and independent role of SMCs in driving arterial disease, but the addition of ECs and circulating immune cells into the disease models would allow for studies aimed at better understanding cell-cell communication and inflammatory activation. At the same time, linking biomolecular changes in the SMCs to tissue-wide dysfunction may help elucidate important molecular targets for therapeutic intervention. Because of this, the ARTOC has the potential to reduce translational failures and enhance human disease research with validated data that can be applied to mechanistic studies, therapeutic screening, and precision medicine across various arterial diseases.

## METHODS

### Fibrin scaffold synthesis and preparation for cell seeding

Fibrin hydrogel scaffolds were fabricated as reported previously ^35^. In short, a solution of bovine fibrinogen (Sigma Aldrich) in 0.2% polyethylene oxide (Sigma Aldrich) was electrospun into a spinning circular bath containing 3 kU Thrombin (BioPharm Laboratories) in 50 mM calcium chloride (Sigma Aldrich). The resulting disc of mechanically stretched and aligned nanofibers was then rolled around 0.6 mm steel PTFE-coated mandrels (Component Supply) to generate a tubular shape by rolling it first with longitudinally aligned fibers, then with circumferentially aligned fibers on top of the first layer. The length of each roll is cut manually and highly tunable to control for overall scaffold thickness. “Thin” scaffolds consist of rolling 1.5 cm of longitudinal fibers and 3 cm of circumferential fibers, and “thick” scaffolds are 3 cm longitudinal and 5 cm circumferential. Hydrogel tubes were further crosslinked overnight in 100 mM NHS/ 60 mM EDC solution and then dehydrated via sequential ethanol baths increaseing in concentration from 25% to 100%, followed by air drying and storage at 4 degrees C for up to 1 month. Prior to seeding cells on the fibrin scaffolds, they were gradually rehydrated in 75% ethanol followed by PBS. They were then coated in a solution of bovine fibronectin (Sigma-Aldrich) and rat tail collagen (Corning) at a concentration of 50 ug/cm^2^ for one hour, followed by overnight incubation in cell culture media at 4C. Tubular fibrin scaffolds were then warmed to 37 degrees C prior to cell seeding.

### Tissue elasticity measured by radial tensile testing

ARTOC were harvested from bioreactors and cut orthogonally into 2 mm long sections, and tissue length and wall thickness were measured under light microscopy. Using an RSA-G2 Mechanical Analyzer, tissues were loaded by inserting two force-transducing pins into the tissue lumen, and the pins were pulled apart at 0.05 mm/sec until total failure. Tension force and gap were measured as a function of time and normalized to sample dimensions using the hoop stress equation to obtain circumferential stress and strain values, and stress vs. strain curves were generated. Stress at failure and strain at failure were obtained from the raw normalized data at the point of ultimate failure. Raw normalized data was fitted to an exponential function to generate fitted stress vs. strain curves, and the fitted curves were used to calculate elastic moduli values for statistical analysis. Incremental elastic moduli were calculated at each point along the stress vs. strain curves by taking the derivative of the fitted exponential function.

### Contractility in response to vasoagonists

For all samples, cells were serum starved by incubation in high glucose DMEM with 1% P/S and 0.5% FBS for 24 hours prior to treatment. For 2D iSMCs, cells culture in a 12 well tissue culture plate were fluorescently stained with CellTracker Green (1 uM) for 1 hour at 37C and acclimated to an incubator module in a Nikon AX-R Confocal Microscope. After a baseline image was taken, wells were treated with 80 mM KCl in the same media and imaged every minute for 5 minutes, followed by every 5 minutes for the remaining 25 minutes. KCl solution was then washed out with media 3x and allowed to incubate for 1 hour, and then wells were treated with sequentially increasing doses of phenylephrine in the same media, from 10^-5^ mM to 10^-1^ mM. PE treatment was imaged in the same way as the KCl treatment, but media was not washed out between each sequential treatment. For ARTOCs, tissues were harvested and dissected into 1 mm thick sections and incubated in a 96 well plate in media, with the cross section facing directly down toward the scope lens. ARTOCs were then treated and imaged with the same method as the 2D iSMCs, only using transmitted light rather than fluorescent light. For both 2D iSMCs and ARTOCs, area was thresholded from the baseline image and quantified for each sequential image, with % contraction defined as: 1 – current area/baseline area. PE % contraction was then normalized to maximum KCl % contraction.

### *In situ* pressure, burst pressure, and distention measurements

Microfluidic pressure transducers (Fluigent Inline Pressure Sensor) were placed directly preceding and following one tubular fibrin scaffold in the bioreactor culture chamber to measure inlet and outlet pressures, respectively. To reduce the difference in resistance between the two tubular fibrin scaffolds of one bioreactor culture chamber, flow resistance was calculated for the inline pressure sensors and used to adjust the length of tubing connected to the other tubular fibrin scaffold. This was done using the Poiseuille equation, with key assumptions that 1) tubing, needles, and the tubular fibrin scaffold can be approximated as rigid cylindrical pipes, and 2) luminal media can be approximated as incompressible and Newtonian:

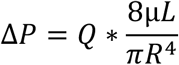

For a range of peristaltic pump flow rates (e.g., 10 – 60 bpm or 0.5 – 3 mL/min), the system was allowed to approach a steady state for 5 minutes before pressure measurements were collected for 2 minutes. Measurements were taken using fibrin-only scaffolds in the bioreactor culture chambers to estimate luminal pressure.

Burst pressure was measured using the same pressure sensor as above, with the outlet capped rather than connected to tubing. A syringe pump was used to steadily flow in dyed media at 0.05 mL/min and rising pressure was recorded in real time until a visible pinhole or total failure was seen by dyed media releasing into the external chamber. This point was time-matched to pressure data and recorded as the burst pressure, followed by conversion to pressure inside the scaffold using the same equation as above.

To measure distention, bioreactor culture chambers were first flushed with phenol red-free SMC media. Bioreactors were then placed vertically in a high-speed optical micrometer (Keyence LS-9000), placed inside the incubator. A laser micrometer (MAKER) was used to measure tubular fibrin scaffold outer diameter at increasing pulsatile flow rates (10 – 60 bpm or 0.5 – 3.0 mL/s). Flow rates were applied for 5 minutes prior to data collection to allow the flow to approach steady state. Distention measurements were collected for 2 minutes for both fibrin scaffolds and ARTOC and averaged between two samples in one bioreactor.

### Simulation of pulsatile radial distenstion and wall shear stress

Simulations were calculated using the open-source program SimVascular (May 2023 release).^39^ Because cell culture media (5% serum) was used to perfuse the vessel in bioreactor experiments, the fluid was modeled as incompressible and Newtonian. The vessel wall was modeled as deformable, with a uniform elastic modulus and uniform thickness along the length. The fluid structure interaction (FSI) between the fluid domain and the vessel wall is simulated using the coupled momentum method (CMM). The fluid solver uses the generalized minimal residual method (GMRES). Inflow boundary conditions were flow rates approximated by 10 Fourier modes, derived from in-line pressure measurements before and after the constructs using a lumped parameter model and assuming Poiseuille flow conditions. Outflow conditions were prescribed boundary resistances based on the lumped parameter model of the flow setup. For computational efficiency, initial conditions for the simulations of pulsatile flow in a deformable vessel were first calculated from simplified simulations using constant flow (the average flow rate from the corresponding pulsatile flow data) in a rigid vessel. Input vascular wall and fluid properties, as well as simulation parameters, are listed below (**Table S1-2**).

**Table S1.** SimVascular Wall and Fluid Properties.

| Parameter | Value |
| --- | --- |
| Wall Type | Deformable, constant |
| Thickness | 0.05 cm |
| Poisson Ratio | 0.5 |
| Elastic Modulus (Cellularized) | 2.25E6 dyne/cm<sup>2</sup> |
| Elastic Modulus (Fibrin Only) | 0.75E6 dyne/cm<sup>2</sup> |
| Density | 1.1 g/cm<sup>3</sup> |
| Shear Constant | 8.33E-1 |
| Fluid Density | 1.002 g/cm<sup>3</sup> |
| Fluid Viscosity | 0.00862 g/cm/s |

**Table S2.** SimVascular Simulation Parameters.

|  |  |
| --- | --- |
| Backflow Stabilization Coefficient | 0.2 |
| Flow Advection Form | Convective |
| Force Calculation Method | Velocity Based |
| Maximum Number of Iterations for svLS Continuity Loop | 400 |
| Maximum Number of Iterations for svLS Momentum Loop | 2 |
| Maximum Number of Iterations for svLS NS Solver | 1 |
| Minimum Required Iterations | 3 |
| Number of Krylov Vectors per GMRES Sweep | 100 |
| Number of Solves per Left-hand-side Formation | 1 |
| Number of Timesteps | 100 |
| Number of Timesteps between Restarts | 10 |
| Output Surface Stress | True |
| Pressure Coupling | Implicit |
| Print Average Solution | True |
| Print Error Indicators | False |
| Quadrature Rule on Boundary | 3 |
| Quadrature Rule on Interior | 2 |
| Residual Control | True |
| Residual Criteria | 0.01 |
| Step Construction | 2 |
| Time Integration Rho Infinity | 0.5 |
| Time Integration Rule | Second Order |
| Time Step Size | 0.001 |
| Tolerance on Continuity Equations | 0.4 |
| Tolerance on Momentum Equations | 0.05 |
| Tolerance on svLS NS Solver | 0.4 |
| svLS Type | GMRES |

### Base editing

The base editor, (BE4max plasmid #112099), was obtained from Addgene. The gRNA (GTGGGCGGACCCATCTCCTC) targeting the LMNA gene (c. 1824 C>T; p.Gly608Gly) at exon 11 was synthesized by Synthego. Human iPSC (SCVI111) were dissociated into single cells using TrypleE express (Thermo), and 250,000 cells were nucleofected (1200V, 20ms, 1 pulse) with 1μg of the BE4max vector and 60 moles of the sgRNA (Synthego), using the Neon Transfection System (Thermo Fisher). Individual iPSC clones were isolated from the Isocell system (Iota Sciences) and expanded. Genomic DNA was extracted from individual clones using Quick Extract solution (Lucigen) and PCR-amplified with GoTaq HotStart polymerase (Promega) (PCR cycling condition: 95 °C 2 min; 95 °C 15sec, 60 °C 15sec, 72 °C 1 min (40 cycles); 72 °C 1 min).The C to T base conversion at LMNA exon 11 of individual clones was verified by Sanger sequencing.

### iPSC expansion and differentiation into iECs and iSMCs (Healthy and HIF2A lines)

iPSCs (C1-2^71^, BC1, and LMNA^WT^, and LMNA^HGPS^ were expanded on tissue culture plastic coated with either Vitronectin, Matrigel or Geltrex in Essential 8 (E8) media supplemented with 10 mM Y-27632 (StemCell Technologies) for the first 24 hours after passaging. iECs were derived using previously published protocols ^50,72^. Briefly, iPSCs on vitronectin were differentiated into iECs by inducing mesoderm lineage in Essential 6 media supplemented with CHIR 99021 (StemCell Technologies) for 2 days, followed by passaging onto collagen-coated plates and cultured in Endothelial Cell Growth Medium (ECGM, PromoCell) supplemented with additional 50 ng/mL VEGF-A (Peprotech) and 50 mM SB431542 (Cayman Chemical) for an additional 6 days, and finally Magnetic-Activated Cell Sorting was used to isolate CD31+ iECs. iECs were expanded up to passage 3 in ECGM supplemented with 25 ng/mL VEGF and 50 mM SB431542 (EC media) <u>on Collagen I-coated plates</u>. A previously established differentiation protocol was used to derive iSMC with minor modifications^73^. In brief, iPSCs cultured on Matrigel underwent mesoderm induction in RPMI medium supplemented with B27 and CHIR 99021 (StemCell Technologies) for 3 days, followed by vascular specification for 4 days in RPMI medium supplemented with 2% B27 without insulin, 25 ng/mL VEGF-A, and 25 ng/mL bFGF (Peprotech). Vascular mural fate was then induced over 7 days in RPMI medium supplemented with B27, 5 ng/mL PDGFb and 2.5 ng/mL TGFb (Peprotech), and the remaining cell population then underwent purification for 7 days in glucose-free RPMI supplemented with insulin-free B27, PDGFb, TGFb, and 4 mM sodium lactate. iSMCs were expanded up to passage 6 in high glucose DMEM supplemented with 5 ng/mL bFGF (Peprotech) and 5% fetal bovine serum (SMC media) on Collagen I-coated plastic. A slightly different protocol was used for LMNA^WT^ and LMNA^HGPS^ cells. In brief, three days before initiating differentiation, the iPSCs were split at a ratio of 1:12. Differentiation was initiated by replacing the E8 medium with DMEM/F-12 (Corning) supplemented with 1X N-2 and 1X B27 supplements (Life Technologies), 6μM CHIR-99021 (Tocris), and 25 ng/ml human BMP4 (Peprotech) (day 1). This medium was maintained for 48 hours without media changes. On day 4, the culture medium was switched to DMEM/F12/N2/B27 with 10 ng/ml PDGF-BB (Peprotech) and 2 ng/ml Activin A (Peprotech). On day 6, cells were dissociated using TrypLE Express (Life technologiesand replated onto gelatin-coated plates (0.1%) in DMEM/F12/N2/B27 medium supplemented with 10 ng/ml PDGF-BB to promote further lineage specification. By day 8, the cells were transitioned to SmGM-2 medium (Lonza) to support growth. Once confluent, the iSMCs were expanded through 2 passages on gelatin-coated plates.

### BrdU cell proliferation assay

After differentiation, iSMCs derived from LMNA^WT^ or LMNA^HGP^) were seeded in 96-well plates. After an additional 72 h, cells were pulsed with BrdU for 3 h before DNA incorporation was measured using the DELFIA Cell Proliferation kit (PerkinElmer, AD0200) according to the manufacturer’s protocol. The fluorescence signal in the 96-well plate was read with an EnVision 2104 plate reader (Perkin Elmer).

### iSMC seeding on fibrin tubular constructs

Following expansion, iSMCs were seeded on 35mm tissue culture dishes at a density of 500,000 per dish in SMC media supplemented with 50 ug/mL L-ascorbic acid (Sigma Aldrich) for 7 days. Then, an additional 250,000 cells per dish were seeded on top of the existing cell layer and cultured for an additional 5-7 days to achieve a multilayer of cells. To initiate the release of iSMCs from the dishes for rolling onto fibrin scaffolds, the dishes were washed with PBS and incubated with a 50 mM L-argenine (Sigma Aldrich) solution in 25 mM HEPES buffer (pH 7.5, Gibco) for 2 minutes at 37 °C. Using a cell scraper to push from the outer edges toward the center of the dish, the iSMCs were gently released from the dish in one contiguous sheet. Using curved forceps, the cell sheet was lifted from the dish and draped over a cannulated tubular fibrin scaffold in an assembled bioreactor (sterilized by autoclave). With 2 pairs of curved forceps, both long ends of the cell sheet were wrapped repeatedly around the scaffold in opposite until the sheet was completely wound in a spiral around the scaffold and allowed to adhere for 30 minutes in a suspended droplet of SMC media (1% Pen/Strep), and then the entire chamber was filled with media and sealed.

### Dynamic flow culture in bioreactors

Once ARTOC were seeded with cells, 0.22um syringe filters were used to cap the bioreactor chamber media tubing to allow for gas exchange. The bioreactor was incubated at 37 degrees C and ARTOC were cultured statically for 1-2 days in SMC Media with 1% Pen/Strep (BR SMC media). The bioreactor was then connected to the flow setup. Peristaltic pump tubing (1/8 inch ID, sterilized with ethanol) was flushed with sterile water, then connected to rest of the reservoir chamber setup (tubing ID 1/16 inch, sterilized by autoclave) and all tubing was flushed with BR SMC media. Tubing was inserted into peristaltic pump (Ismatec IP-C), which was turned on to pump any remaining air bubbles out of the tubing. The tubing was connected to the bioreactor culture chamber and 0.22um syringe filters were installed on the reservoir chambers to enable luminal gas exchange. The entire setup was returned to the incubator. ARTOC were cultured with luminal flow of BR SMC media at 12 bpm (0.5 mLmin) for up to 3 weeks. Bioreactor culture chamber and reservoir chambers were refreshed with BR SMC media every three days.

### iEC seeding and permeability assay

Rehydrated fibrin scaffolds were sutured on both ends, with one end tied tight and the other left loosely around the circumference of the scaffold. Tubular fibrin scaffolds were transferred to microfuge tubes and stood vertically. iECs were suspended in EC media with 1% Penicillin/Streptomycin (pen/strep, Gibco) at 40 million cells/mL. Using a sterile syringe and 27G blunt tip hollow needle, 20 uL of the cell suspension was injected into the scaffold through the loose end, and then the loose suture is tied tight and the microfuge tube was filled with EC media. Tubes were place horizontally in incubator at 37 degrees C and rotated manually 90 degrees every 10 minutes for 80 minutes. After rotation was completed, tubes were opened slightly and incubated vertically for 2 days, with media changed after 24 hours. To test iEC permeability in ARTOC, seeded scaffolds were cultured in EC media with 1% pen/strep for 4 days prior to assay. Two seeded scaffolds and two fibrin-only scaffolds were cannulated into bioreactor culture chambers with ECGM media supplemented with 1% pen/strep, and luminal flow was started immediately with the same media supplemented with 0.25 mg/mL FITC-dextran (55kDa-77kDa, Sigma-Aldrich). Aliquots of both luminal and chamber media were taken immediately upon flow startup and at the 24 hour point after flow began. All aliquots were measured for fluorescent intensity at 528 nm using a plate reader, and permeability coefficient was calculated using a previously established endothelial permeability equation ^74^. Seeded scaffolds were then imaged to confirm iEC monolayer presence and quantify retention of FITC-dextran within the scaffold itself.

### Immunostaining and image quantification

For LMNA WT and HGPS lines in 2D culture, the iSMCs were fixed with 4% PFA for 10 min at room temperature, washed 3 times with DPBS, followed by permeabilization with 0.1% Triton for 10 min at room temperature. After blocking for 1 h at room temperature with blocking buffer (DPBS/2%FBS/2% BSA), the iSCMs were incubated with primary antibodies, Lamin A/C (1:500, Bio-rad, MCA1429GA, clone JOL2) and Smooth muscle actin (1:400, Cell signaling #19245) at 4 °C overnight. The cells were washed 3 times for 5 min each with DPBS and incubated with Alexa-conjugated secondary antibodies for 1 h at room temperature. After washing 3 times for 5 min each, a drop of NucBlue was added to counterstain the DNA. Immunofluorescence Images were obtained using a confocal microscope (Zeiss LSM880).

For ARTOC staining, tissues were harvested from the bioreactor culture chambers, fixed in 10% formalin (Sigma-Aldrich) at 4 degrees C overnight, washed with PBS, and then dehydrated in 20% sucrose overnight at 4C. ARTOC were frozen in optimal cutting temperature compound (OCT) and cryosectioned axially at a thickness of 10 microns. Once on slides, sections are blocked in a solution of 5% bovine serum albumin (BSA, Sigma-Aldrich), 0.5% Triton-X (Alfa-Aesar), and 0.05% Tween20 (Sigma-Aldrich) in PBS for an hour at room temperature. Primary antibodies were diluted in the same blocking solution and incubated at 4 degrees Celsius overnight. Primary antibody solution was removed and samples are washed with PBS + 0.05% Tween-20. Secondary antibodies were added at a 1:400 dilution in the blocking solution and incubated at room temperature for 2 hours before washes with PBS + 0.05% Tween20. DAPI was added at a 1:1000 dilution and incubated at room temperature for 10 minutes and washed off with PBS. The following primary antibodies were used: mouse anti-VE-cadherin (BBA19, 1:200), mouse anti-aSMA (AB7817, 1:100), rabbit anti-smooth muscle myosin heavy chain (SM-MHC, AB82541, 1:200), rabbit anti-collagen I (AB34710, 1:200), mouse anti-collagen III (AB6310, 1:200), mouse anti-elastin (AB77804 1:50), rabbit anti-fibronectin (AB32419, 1:200), rabbit anti-p21 (CST2947S, 1:200), mouse anti-progerin (sc-81611, 1:100) and phalloidin-AlexaFluor 546 (Invitrogen, 1:400). Sections were imaged using the Nikon AX-R inverted confocal microscope. Images were quantified in FIJI/ ImageJ. For each section, a max intensity Z projection was taken. The fibrin was then masked out and the tissue area is measured. Nuclei were counted using auto-thresholding method (Otsu) from the DAPI channel. For each channel, a threshold for intensity was determined manually and these values were set to create a mask which was applied batchwise to all samples being compared. Mean intensity and area of the mask was recorded for quantification of area and intensity of protein expression. Representative images shown were processed as follows: background was subtracted using a rolling ball correction, despeckling with a radius of 1 pixel was applied, brightness was increased by 40% and a 20% contrast was applied on all images to visualize protein expression clearly.

### Atomic force microscopy

ARTOCs were isolated and immediately flash frozen in liquid nitrogen. Frozen grafts were embedded (Tissue-Tek OCT Compound) and left at −80C overnight. Frozen samples were then cryosectioned (epredia HM525 NX) in 30 µm slices before being adhered to positively charged glass slides (fisherbrand 1255015). Tissues were submerged in 1x protease and phosphatase inhibitor (Thermo Fisher Scientific 78442) in PBS before measurement to prevent degradation. AFM measurements were performed while submerged using a Nanowizard V BioScience (Bruker) equipped with SAA-SPH-1UM probes with a setpoint of 3 nN. Indentation tests were performed in iSMC regions. Regions of size 25 by 25 µm were selected, with 25 measurements taken in the selected region. The probe oscillated in frequency shuffle of 1, 2.2, 4.5, 10, 22, 45, and 100 Hz for at least 5 cycles with an amplitude of 20 nm. The Storage Modulus, Loss Modulus, and Loss Factor were calculated by fitting the force-displacement oscillation curves with an assumed Poisson’s ratio of 0.5, with poorly fitting curves filtered out.

### Statistics

Biological replicates are indicated as N and technical replicates are indicated as n. Biological replicates are defined as separate differentiation replicates from the parental iPSC lines, and technical replicates come from 2 ARTOCs from the same N cultured in the same bioreactor chamber at once. For all statistical tests, n’s were first averaged for each N, and tests were run solely on the N’s. Details of replicates for each experiment can be found in the figure legends. All bar graphs represent means ± SEM. Graphpad Prism (v10.2.3.) was utilized for statistical analysis. Ratio paired t-test was used for wall thickness, tensile testing, immunofluorescent imaging quantification, permeability, elastin content, and AFM-microrheoology measurements. This method was used for these experiments to correct for biological variability by directly comparing between ARTOCs of the same biological replicate, or to directly compare between fibrin vs. ARTOC evaluated in parallel in the same experimental replicate. Unpaired Welch’s t-test was used for burst pressure, distention, WSS, and vasoagonist response measurements between fibrin constructs or 2D iSMCs and ARTOCs with significantly different standard deviation (ratio of 2 or greater) between groups, while a standard unpaired t-test was employed for the same comparisons with similar standard deviation (ratio less than 2) between groups. This method was used when fibrin and cellularized samples required evaluation at separate experimental timepoints so that pairing was not relevant. Lastly, a 1-sample t-test was used to compare each simulated distention value as a mean value to each experimental distention group because simulations used the same values within each comparison did not have experimental replicates.

## ACKNOWLEDGEMENTS

We would like to thank Dr. Chelsey Dunham for bioreactor chamber design and training on scaffold fabrication, Dr. Franklyn Hall for training on SMC differentiation and seeding, and Marcos Negrete and Makenzie Bushold for technical support. DY and CA are supported by the NSF-GRFP. We acknowledge support from the Duke Cancer Institute as part of the P30 Cancer Center Support Grant (Grant ID: P30 CA014236). This work was supported by RAD0102 and SNT0101 from the Translational Research Institute through NASA Cooperative Agreement NNX16AO69A (both to SG).

## SUPPLEMENTARY FIGURES

**Figure S1.**
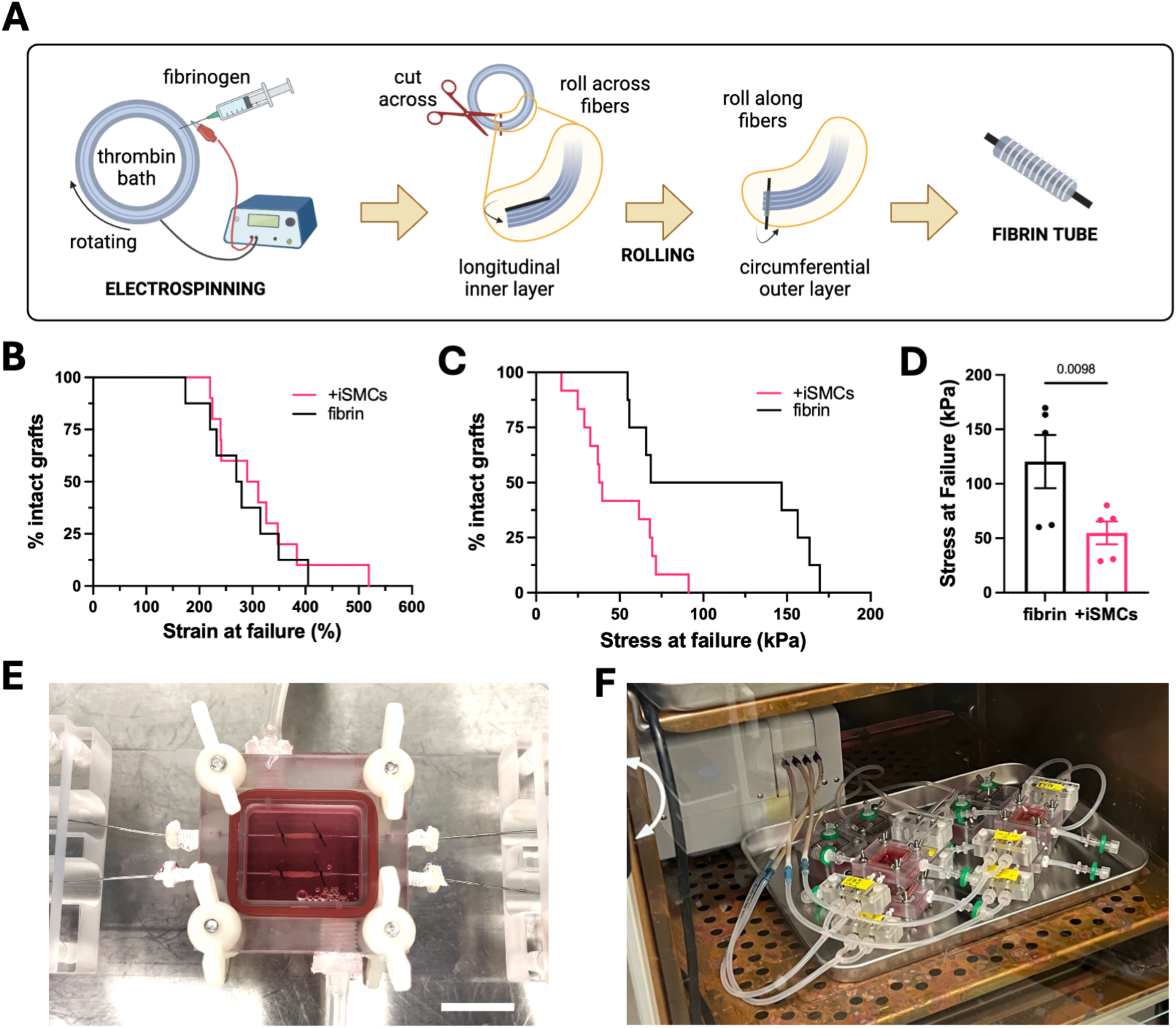
Artery-on-chip characterization and culture. **(A)** schematic of fibrin hydrogel electrospinning and rolling to generate a tubular construct. Survival curves for ultimate failure of fibrin vs. ARTOC with **(B)** increasing strain and **(C)** increasing stress at failure. **(D)** Ultimate tensile stress at total failure in fibrin vs. ARTOC, N=5 (n=2-3 each N). **(E)** image of cellularized ARTOC in a bioreactor chamber (scale 1 cm) and **(F)** pulsatile flow culture system connected to a peristaltic pump.

**Figure S2.**
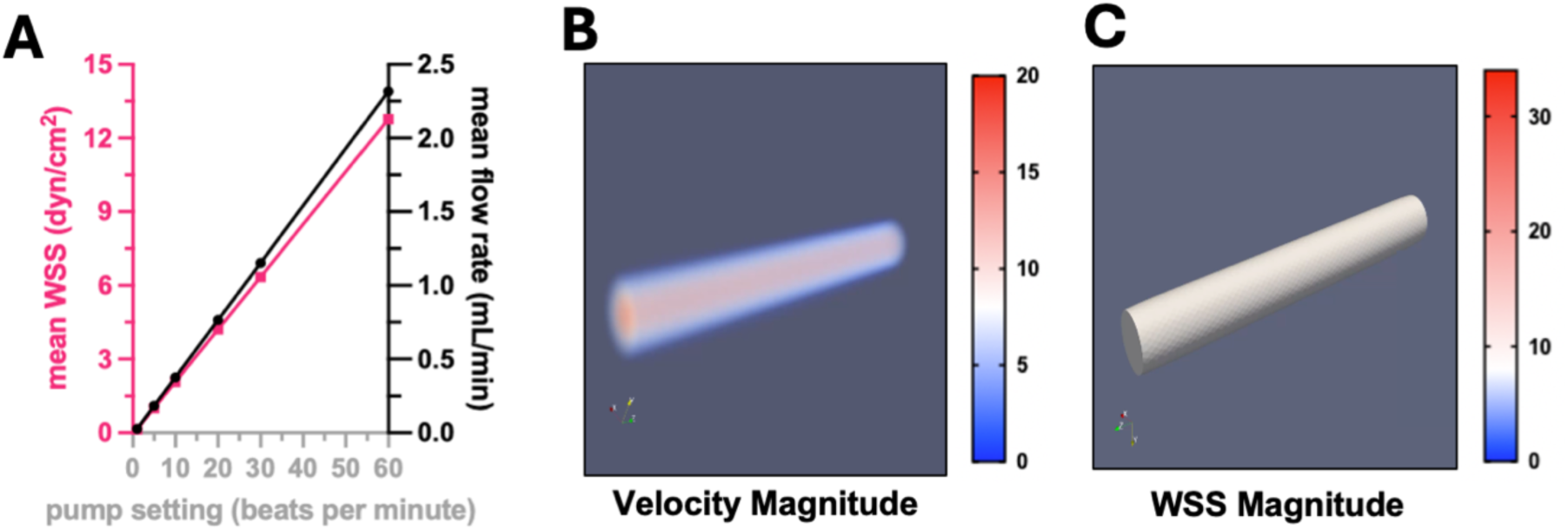
(A) Measured flow rate (right axis) and estimated mean WSS calculated via Hagen-Poiseuille assumptions (left axis) corresponding to pump speed. (B) snapshot of simulated luminal velocity profile over one time loop (6 seconds) at 0.5 mL/min. See Video S1(C) snapshot of simulated WSS profile over one time loop (6 seconds) at 0.5 mL/min. See Video S2

**Figure S3.**
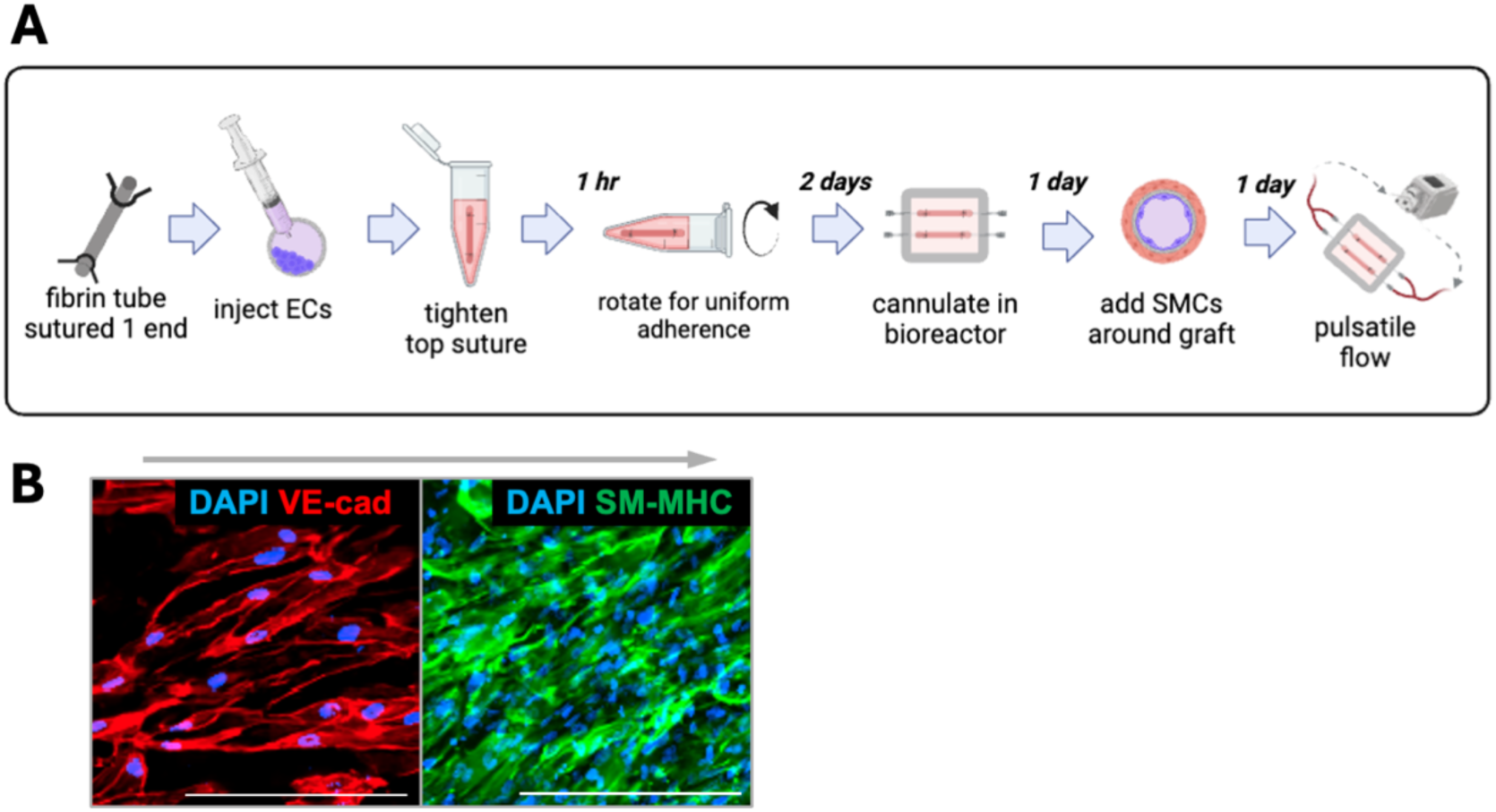
Co-culture ARTOC assembly and culture. (A) schematic for EC seeding protocol and co-culture ARTOC assembly. (B) en face staining of co-culture ARTOC on luminal side (left) for VE-cad in iECs and exterior (right) for SM-MHC for iSMCs, gray arrow indicates direction of flow, scale=50 um.

**Figure S4.**
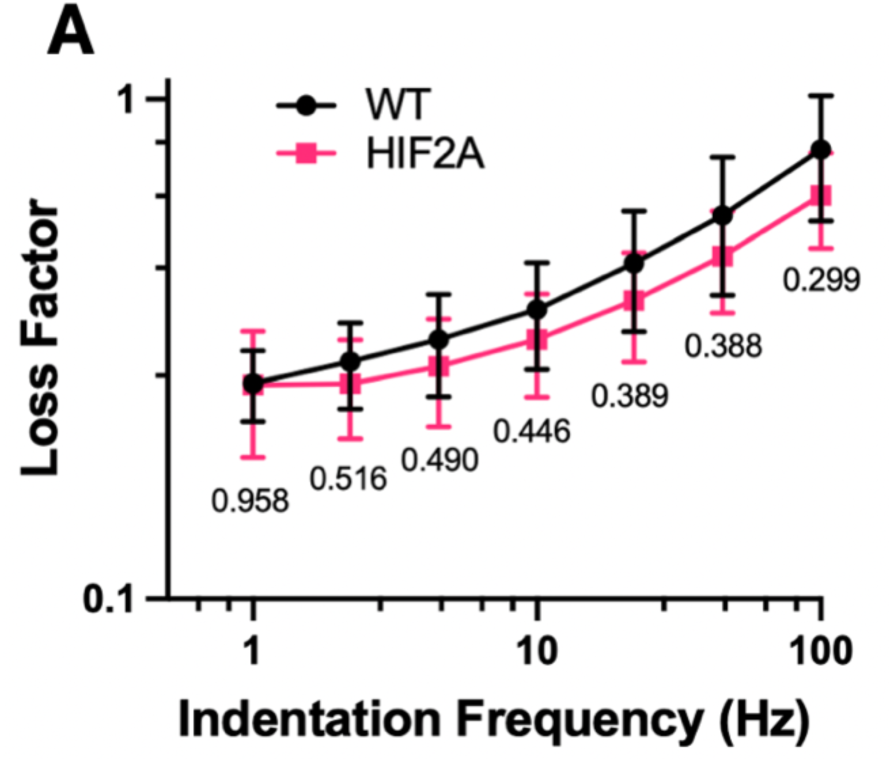
HIF2A Microrheology in ARTOCs. (A) loss factor ratio between elastic and viscous properties in HIF2A or WT ARTOC measured over an entire 1 to 100 Hz frequency sweep, N=4 (n=65-130 each N)

**Figure S5.**
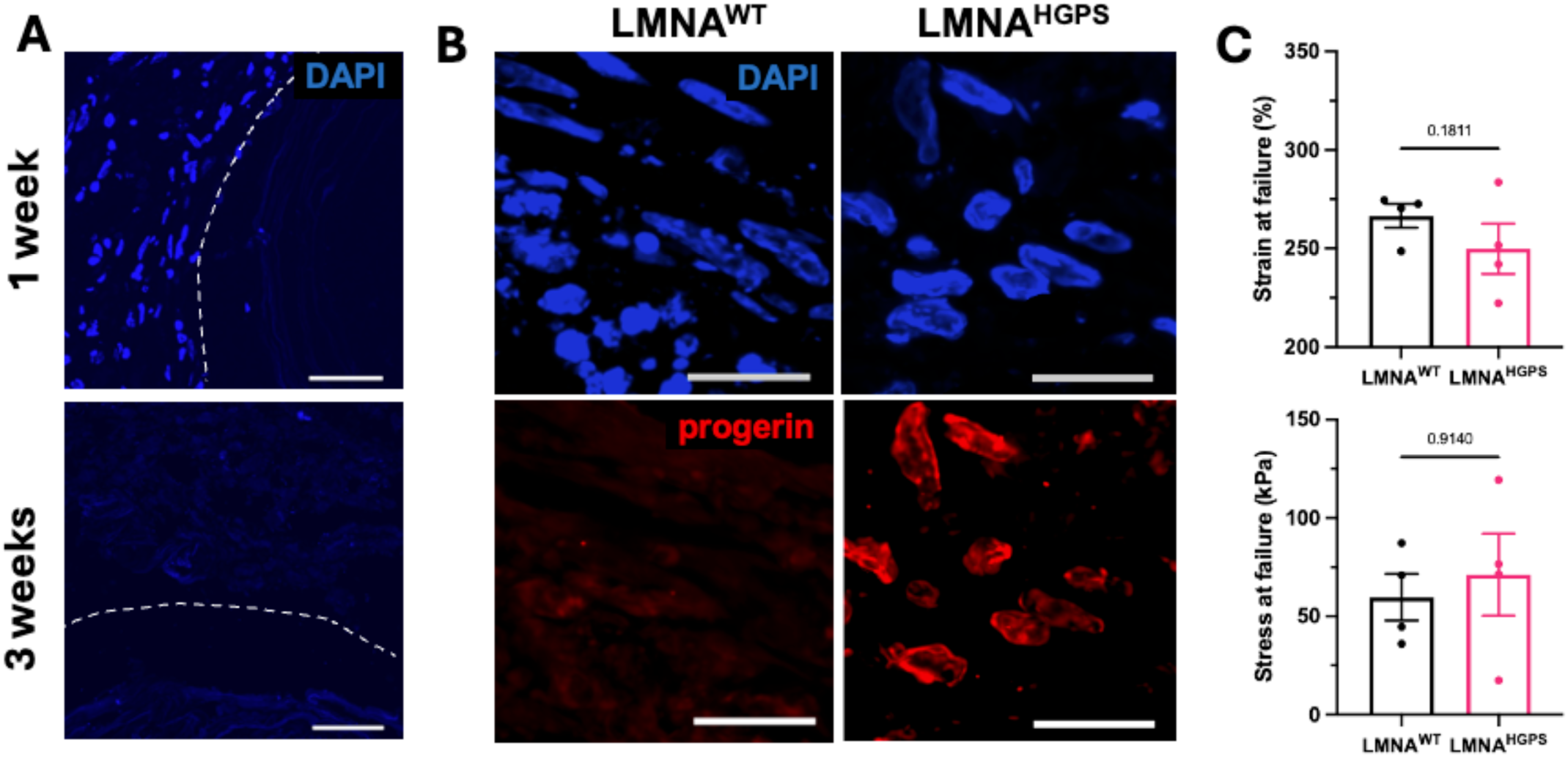
Further characterization of LMNA^HGPS^ ARTOC phenotype. (A) nuclear staining for LMNA^HGPS^ after 1 (top) or 3 (bottom) weeks of culture time, dotted line=lumen, scale=50um. (B) Cross-sectional staining in LMNA^WT^ and LMNA^HGPS^ ARTOC for progerin (red) after 1 week of culture under flow, scale=20um. (C) strain (top) and stress (bottom) at ultimate failure in LMNA^WT^ and LMNA^HGPS^ ARTOC after 2 weeks of culture, N=4 (n=2-3 each N).

## Notes

### Competing Interest Statement

The authors have declared no competing interest.

